# PARP12 is required to repress the replication of a Mac1 mutant coronavirus in a cell and tissue specific manner

**DOI:** 10.1101/2023.06.16.545351

**Authors:** Catherine M. Kerr, Srivatsan Parthasarathy, Nancy Schwarting, Joseph J. O’Connor, Emily Giri, Sunil More, Robin C. Orozco, Anthony R. Fehr

**Affiliations:** Department of Molecular Biosciences, University of Kansas, Lawrence, Kansas 66045, USA; Department of Veterinary Pathology, Oklahoma State University, Stillwater Oklahoma 74048, USA

**Author notes:** Correspondence; (K.S.).

**Keywords:** Coronavirus, MHV, ADP-ribosylation, PARP, macrodomain, pathology

## Abstract

ADP-ribosyltransferases (ARTs) mediate the transfer of ADP-ribose from NAD^+^ to protein or nucleic acid substrates. This modification can be removed by several different types of proteins, including macrodomains. Several ARTs, also known as PARPs, are stimulated by interferon, indicating ADP-ribosylation is an important aspect of the innate immune response. All coronaviruses (CoVs) encode for a highly conserved macrodomain (Mac1) that is critical for CoVs to replicate and cause disease, indicating that ADP-ribosylation can effectively control coronavirus infection. Our siRNA screen indicated that PARP12 might inhibit the replication of a MHV Mac1 mutant virus in bone-marrow derived macrophages (BMDMs). To conclusively demonstrate that PARP12 is a key mediator of the antiviral response to CoVs both in cell culture and *in vivo*, we produced PARP12^−/−^ mice and tested the ability of MHV A59 (hepatotropic/neurotropic) and JHM (neurotropic) Mac1 mutant viruses to replicate and cause disease in these mice. Notably, in the absence of PARP12, Mac1 mutant replication was increased in BMDMs and in mice. In addition, liver pathology was also increased in A59 infected mice. However, the PARP12 knockout did not restore Mac1 mutant virus replication to WT virus levels in all cell or tissue types and did not significantly increase the lethality of Mac1 mutant viruses. These results demonstrate that while PARP12 inhibits MHV Mac1 mutant virus infection, additional PARPs or innate immune factors must contribute to the extreme attenuation of this virus in mice.

**IMPORTANCE:** Over the last decade, the importance of ADP-ribosyltransferases (ARTs), also known as PARPs, in the antiviral response has gained increased significance as several were shown to either restrict virus replication or impact innate immune responses. However, there are few studies showing ART-mediated inhibition of virus replication or pathogenesis in animal models. We found that the CoV macrodomain (Mac1) was required to prevent ART-mediated inhibition of virus replication in cell culture. Here, using knockout mice, we found that PARP12, an interferon-stimulated ART, was required to repress the replication of a Mac1 mutant CoV both in cell culture and in mice, demonstrating that PARP12 represses coronavirus replication. However, the deletion of PARP12 did not fully rescue Mac1 mutant virus replication or pathogenesis, indicating that multiple PARPs function to counter coronavirus infection.

## INTRODUCTION

Coronaviruses (CoVs) are the most prominent viruses in the *Nidovirales* order. CoVs are large positive-sense RNA viruses that cause significant human and veterinary diseases and have been responsible for several outbreaks of lethal human disease in the past few decades, including SARS-CoV-1 and MERS-CoV, which emerged in 2002 and 2012, respectively. In December 2019, a new human CoV emerged from China, SARS-CoV-2, causing Coronavirus Disease 2019 (COVID-19) (1). In March of 2020, the World Health Organization declared the SARS-CoV-2 outbreak as a pandemic, making it the first pandemic to be caused by a CoV (1).

CoVs have a 30 kb genome, which encodes for 20-30 proteins (2, 3). There are four main structural proteins, spike (S), envelope (E), membrane (M), and nucleocapsid (N). CoVs can have up to nine accessory proteins, which are unique to each CoV lineage and are important for the evasion of the immune system (4). There are also sixteen non-structural proteins (nsps) required for virus replication (3). In addition to their roles in viral replication, several nsps also have a role in the evasion of the innate immune response. For example, non-structural protein 3 (nsp3) of coronaviruses encodes for a highly conserved macrodomain, termed Mac1, that has ADP-ribose binding and ADP-ribosylhydrolase (ARH) activity. These activities are conserved across the Hepeviridae, Togaviridae, and Coronaviridae families (5, 6).

Macrodomains are well-described structural domains of ~20 kDa with central β-sheets flanked by α-helices (7). The macrodomains from each of these viral families can promote virus replication or pathogenesis (8). The CoV Mac1 also counteracts the host immune response, as mutation of a highly conserved asparagine shown to ablate ARH activity leads to increased levels of IFN-I and other cytokines (9, 10). Using Murine Hepatitis Virus (MHV) strain JHM (JHMV) as a model, we previously showed that this asparagine-alanine mutation (N1347A) led to decreased virus replication in Type I Interferon (IFN-I) competent, but not IFN-I null cells. Notably, these Mac1-deficient viruses are extremely attenuated *in vivo*, causing little to no disease compared to a wildtype (WT) CoVs in several lethal models of CoV infection (11).

ADP-ribosylation is a common, reversible post-translational modification, defined as the addition of ADP-ribose units onto target protein or nucleic acid. It is known to affect a variety of cellular processes, including cell signaling, DNA repair, and apoptosis (12, 13). Also, it is crucial for the host response to virus infections and several other stress responses. In addition, many bacterial toxins utilize ADP-ribosylation to shut down host processes (14). This modification can contain one or more consecutive ADP-ribose units, resulting in either mono- or poly-ADP-ribosylation (MAR and PAR). Both MAR and PAR are carried out by ADP-ribosyltransferases (ARTs) which utilize NAD^+^ as a substrate to MARylate or PARylate target proteins (15). These ARTs include diphtheria toxin-like (ARTD) and cholera-toxin like (ARTC) families, of which ARTDs carry out most of the ADP-ribosylation in mammalian cells. The ARTDs were formerly known as PARPs, though individual ARTDs are still known by their PARP nomenclature (i.e. PARP1, PARP2, etc.) (16). There are 17 mammalian PARPs, and several are interferon-stimulated genes (ISGs). PARPs can have both pro- and anti-viral effects (17, 18). The PARPs with pro-viral activity include PARP1, PARP7, and PARP11, which can reduce IFN-I production or IFN-I signaling, leading to the enhancement of virus replication (19–22). The PARPs with antiviral activities include PARP7, PARP9, PARP10, PARP11, PARP12, and PARP13 (17).

PARP12 is a mono-ADP-ribosyltransferase that inhibits the replication of several viruses. It can mildly inhibit vesicular stomatitis virus (VSV) when overexpressed in HEK293T cells (23). The over expression of PARP12 from a Venezuelan equine encephalitis virus (VEEV) vector strongly restricted the replication of several other viruses, including Sindbis virus (SINV), encephalomyocarditis virus (EMCV), and VEEV (18). PARP12 was also identified in a screen for ISGs that inhibit the Zika virus (ZIKV) replication (24). Further results showed that PARP12, in coordination with PARP11, was required for the ADP-ribosylation, ubiquitination, and degradation of two ZIKV proteins involved in virus replication (25). Also, PARP12 has also been shown to bind to TRIF and enhance NF-κB activation, which indicates that it may have a role in the inflammatory response (26).

Recently, using PARP inhibitors and NAD^+^ boosters, we demonstrated that ADP-ribosylation was responsible for the attenuation of MHV-JHM (JHMV) N1347A replication and enhanced the IFN-I response to this virus, but had no impact on WT virus (10, 27). These results supported the hypothesis that 1 or more PARPs are potent inhibitors of CoV replication and that CoVs have evolved to encode a protein that is specifically required to antagonize them. To identify the specific PARP(s) that restrict CoV replication, we performed an siRNA screen in primary bone-marrow derived macrophages (BMDMs) targeting most of the IFN-induced PARPs. From this screen we found that knockdown of PARP12 and PARP14 enhanced the replication of N1347A. Knockdown of PARP12 demonstrated the greatest enhancement of N1347A replication, indicating that it may be an inhibitor of CoV replication (10). Notably, a separate study found that PARP12 bound to SARS-CoV-2 RNA and that knockdown of PARP12 in Calu-3 cells enhanced SARS-CoV-2 genomic RNA production, providing further evidence that PARP12 may be a potent inhibitor of CoV replication (28). While powerful, there are some limitations of RNAi knockdown, most notably that a portion of the protein is still present and that there is the potential for off-target effects. To fully understand the role of PARP12 on CoV replication, complete deletion models are necessary.

Here, we created PARP12^−/−^ mice to further explore the role of PARP12 in MHV replication both in cell culture and *in vivo*. We found that the deletion of PARP12 fully rescued Mac1 mutant replication in primary macrophages but did not enhance viral replication in dendritic cells. Furthermore, Mac1 mutant replication and pathogenesis was partially restored in PARP12^−/−^ mice, following JHMV intracranial infection or A59 liver infection. These results indicate that PARP12 contributes to the antiviral response to CoV infection but also that other PARPs must function during infection to prevent Mac1 mutant virus from causing severe disease *in vivo*.

## RESULTS

### Generation of PARP12^−/−^ mice

To test the hypothesis that PARP12 is a host factor capable of inhibiting CoV replication, we generated a PARP12 knockout (KO) mouse to test its role both in cell culture and *in vivo*. The PARP12 KO was engineered by replacing PARP12 with lacZ (for details see Methods) (Fig. 1A). To determine the genotypes of the mice, primers were used that separately detect the presence of the WT PARP12 gene and the lacZ insert (Fig. 1A, Methods). We tracked the overall number of PARP12^+/+^, PARP12^+/−^, and PARP12^−/−^ mice born from PARP12^+/−^ × PARP12^+/−^ crosses over the course of a full year, and we found that WT, Het, and KO mice were born at the expected Mendelian ratios (Fig. 1B). We measured PARP12 expression from each organ using qPCR analysis (Fig. 1C). PARP12 was expressed in both PARP12^+/+^ and PARP12^+/−^ in all organs, while it was completely absent in PARP12^−/−^ mice as expected, confirming that we had knocked out PARP12 (Fig. 1C). PARP12 expression was highest in the heart, lung, liver, and testes (male mice), and lowest in the brain and spleen (Fig. 1C). To determine if the PARP12^−/−^ mice developed normally, several organs including the brain, heart, lungs, liver, kidneys, spleen, and testes from PARP12^+/+^, PARP12^+/−^, and PARP12^−/−^ mice were harvested and weighed (Fig. 1D-E). The organ weights from the PARP12^−/−^ mice were largely comparable to those from the PARP12^+/+^ mice and the PARP12^+/−^ mice, indicating normal development of PARP12^−/−^ mice (Fig. 1-D-E). However, the testes of male mice were smaller than WT mice, though this difference was not statistically significant. Finally, to determine if the loss of PARP12 impacted the development of immune cells, we collected cells from the spleen and analyzed the frequency of innate and adaptive immune cells in 14 week-old PARP12^+/+^ and PARP12^−/−^ mice. We found that PARP12 KO had no impact on the percentages of these cells in the spleen as the frequency of innate immune cells such as macrophages and DCs, and adaptive immune cells such as CD8 T cells, CD4 T cells, and B cells were all nearly identical between PARP12^+/+^ and PARP12^−/−^ mice (Fig. 1F-G, Fig. S1).

**Figure 1.**
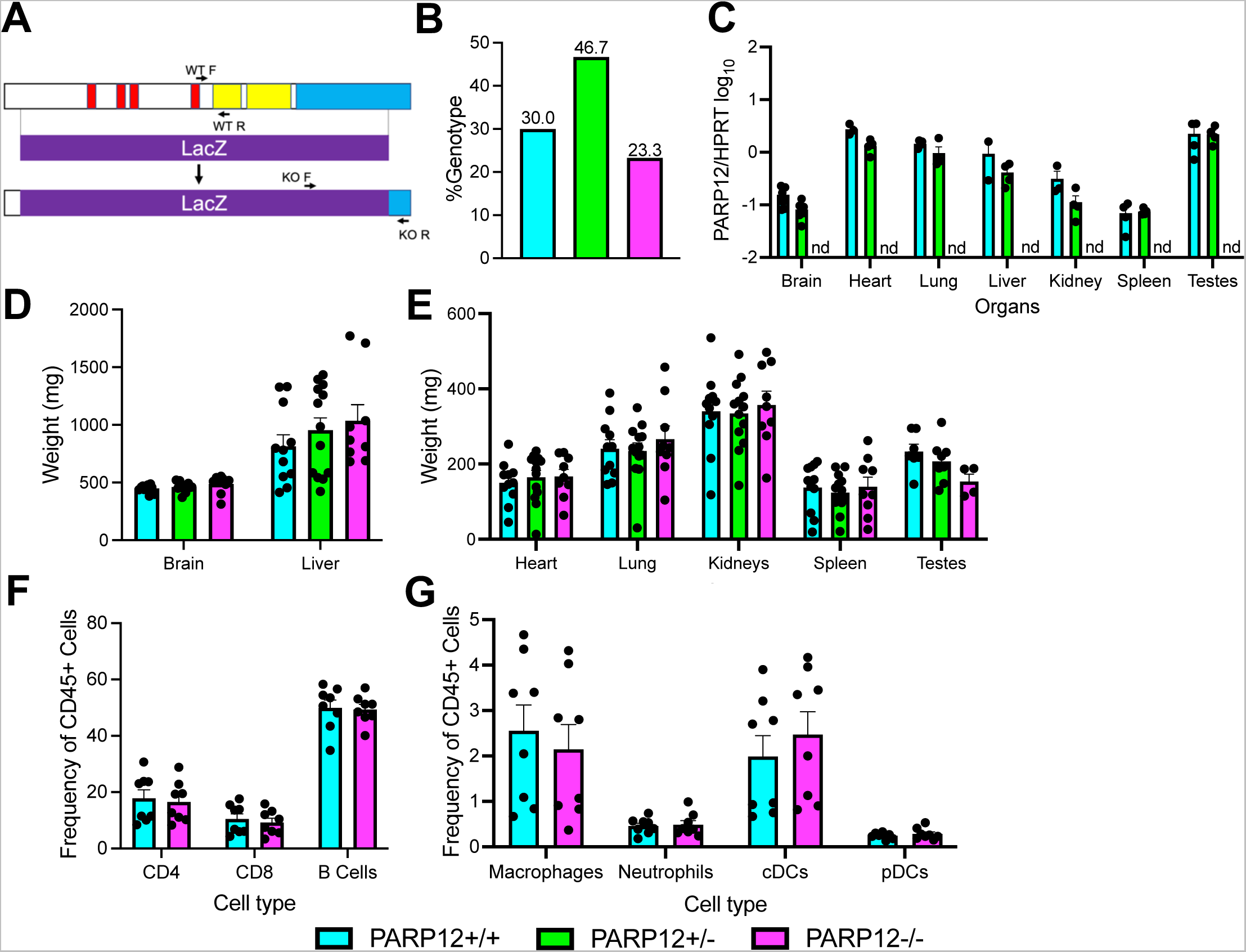
Generation of PARP12^−/−^ mice. (A) Schematic of the LacZ insertion used to create the PARP12 knockout in C57B6/NJ mice. The insertion induces a frameshift mutation, creating a completely null mutation. (B) Ratio of PARP12^+/+^, PARP12^+/−^, and PARP12^−/−^ mice following PARP12^+/−^ × PARP12^+/−^ breeding over the course of one year (n=3). (D-E) PARP12 expression (D) and weights (E) in various organs of PARP12^+/+^, PARP12^+/−^, and PARP12^−/−^ mice (n=4). (F-G) Immune cells from the spleens of naïve PARP12^+/+^ mice and PARP12^−/−^ mice (n=8).

One notable issue we identified with the PARP12^−/−^ mice was that we were unable to produce litters from a PARP12^−/−^ × PARP12^−/−^ pairing. While surprising, this is not without precedent, as several PARPs have been shown to impact the reproductive system (29–32). To formally test this observation, we allowed 3 breeder pairs of PARP12^+/+^, PARP12^+/−^, and PARP12^−/−^ mice to breed over the course of 4 months. We found that PARP12^+/+^ breeders produced 6 successful pregnancies, PARP12^+/−^ mice produced 4 successful pregnancies, while PARP12^−/−^ mice failed to become pregnant (Table S1). However, in rare occasions we were able to get a PARP12^−/−^ female pregnant when bred with a PARP12^+/−^ male, but not vice versa. These results demonstrate that PARP12^−/−^ mice have some defect in their ability to establish pregnancy, which could be tied to the reduced size of the testes in PARP12^−/−^ male mice.

### PARP12 is required for the restriction of Mac1-mutant virus replication in BMDMs

We previously found that siRNA knockdown of PARP12 enhanced, but did not fully restore, the replication of a JHMV N1347A in bone-marrow derived macrophages (BMDMs) (10). Due to the limitations of siRNA knockdown, we hypothesized that a greater enhancement of N1347A replication would be observed with PARP12 knockout cells. We harvested bone marrow cells from PARP12^+/+^ and PARP12^−/−^ mice and then differentiated them into macrophages in cell culture (M0). We then infected the BMDMs with JHMV WT and N1347A at an MOI of 0.05 PFU/cell and collected cells and cell supernatants at 12, 20, and 24 hpi (Fig. 2A). While N1347A replicated at significantly lower levels than WT virus in PARP12^+/+^ cells at 12 and 20 hpi, it replicated at WT virus levels in PARP12^−/−^ cells, demonstrating that PARP12 is required to inhibit JHMV N1347A replication. Importantly, there was no difference in WT virus replication between PARP12^+/+^ and PARP12^−/−^ BMDMs, indicating that PARP12 only inhibits JHMV replication in the absence of Mac1 activity and that Mac1 counters PARP12 activity in these cells. Interestingly, we found that N1347A replication reached near WT levels in PARP12^+/+^ cells at 24 hpi, which we hypothesize was due to a reduction in PARP12 activity in the later stages of infection because of the depletion of NAD^+^ during infection (27).

**Figure 2:**
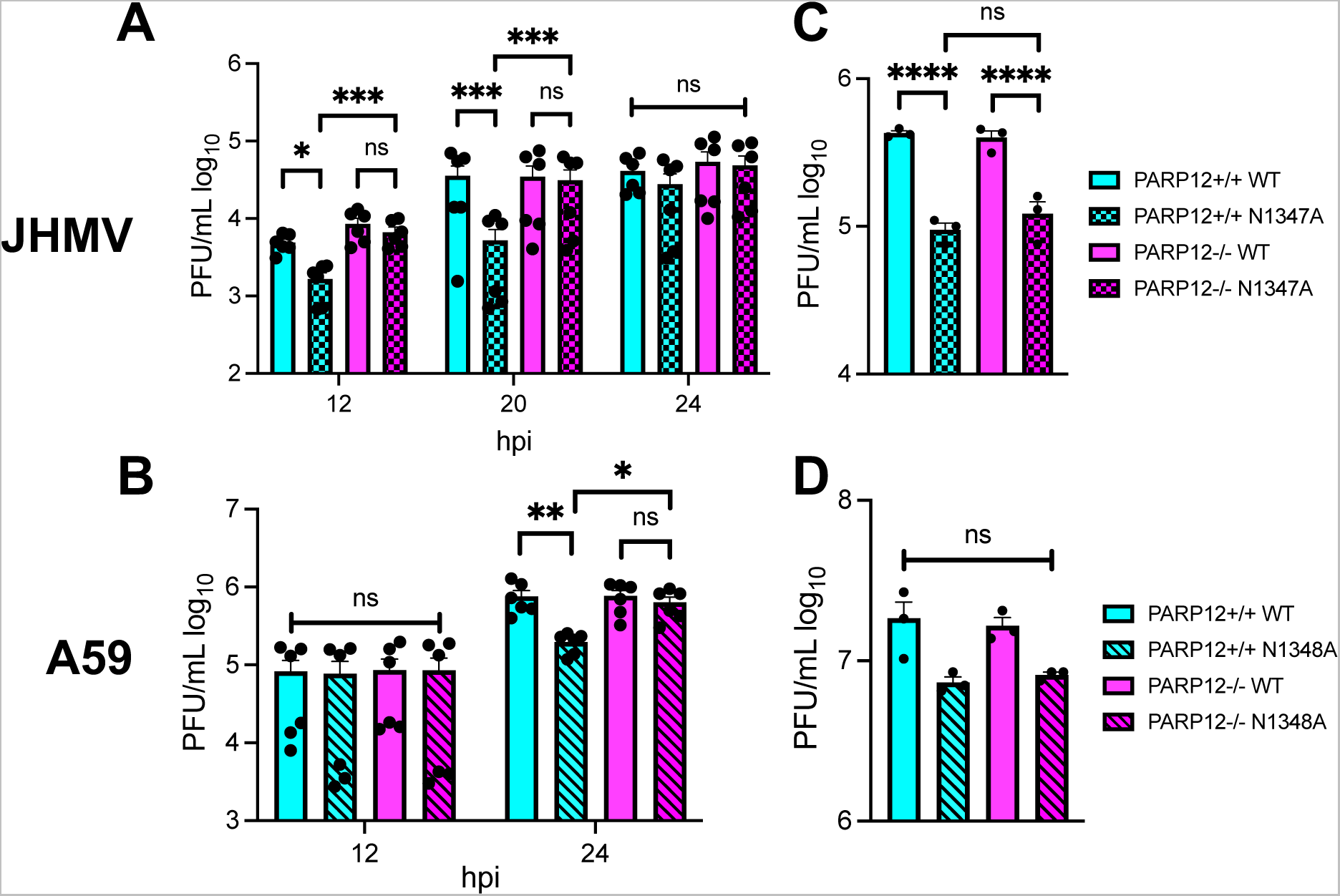
PARP12 is required for the restriction of Mac1-mutant MHV replication in BMDMs, but not BMDCs. (A-B) PARP12^+/+^ and PARP12^−/−^ BMDMs were infected with MHV JHM WT or N1347A (A) or MHV-A59 WT and N1348A (B) at an MOI of 0.05 PFU/cell. Cells and supernatants were collected at indicated times post-infection (hpi) and assayed for progeny infectious virus by plaque assay. The data in A-B are the combined results of 2 independent experiments (n=6). (C-D) PARP12^+/+^ and PARP12^−/−^ BMDCs were infected with MHV JHM WT or N1347A (C) or MHV-A59 WT and N1348A (D) at an MOI of 0.05 PFU/cell. Cells and supernatants were collected at indicated times post-infection (hpi) and assayed for progeny infectious virus by plaque assay. The data in C-D is from one experiment representative of 3 independent experiments (n=3).

To further expand our results to additional strains of MHV, we tested the replication of the MHV-A59 Mac1 mutant virus, N1348A (equivalent mutation to N1347A in JHMV), in both PARP12^+/+^ and PARP12^−/−^ BMDMs. At 12 hpi, there was no difference in replication between WT and N1348A in both PARP12^+/+^ and PARP12^−/−^ BMDMs. But by 24 hpi, there was a significant reduction in the replication of the N1348A virus in the PARP12^+/+^ BMDMs compared to WT virus. In contrast, no difference was detected between N1348A and WT virus replication in PARP12^−/−^ cells (Fig. 2B). This confirmed that PARP12 is necessary for the restriction of the N1348A virus replication in BMDMs.

Next, we tested whether PARP12 was required for the inhibition of JHMV N1347A in other myeloid derived cells. Bone marrow cells were harvested from PARP12^+/+^ mice and PARP12^−/−^ mice and were differentiated into dendritic cells using GM-CSF. Similar to macrophages, N1347A had a significant replication defect in these cells of ~10-fold at 20 hpi. Remarkably, unlike macrophages, the replication defect was not rescued or even enhanced in the PARP12^−/−^ cells (Fig. 2C and D). These results indicate that PARP12 may function in a cell type-specific manner to repress N1347A replication, and that other PARPs are capable of restricting N1347A in GM-CSF derived DCs.

Recently we identified another JHMV Mac1 point mutant that was highly attenuated in cell culture. It was significantly more attenuated than N1347A across multiple cell lines, indicating that Mac1 may have multiple functions during the viral lifecycle (33). This mutant virus (D1329A) was rescued by PARP inhibitors and further inhibited by addition of nicotinamide riboside (NR), clearly demonstrating that it is restricted by ADP-ribosylation. To determine if PARP12 also inhibited the replication of D1329A we infected BMDMs with WT and D1329A viruses and analyzed their replication in PARP12^+/+^ and PARP12^−/−^ cells as described above. In contrast to N1347A, D1329A was not rescued in PARP12^−/−^ BMDMs (Fig. S2). These results again indicate that multiple PARPs are capable of inhibiting CoV replication.

### PARP12 does not impact the IFN-I response during an N1347A infection

Previously we found that N1347A induces an increased IFN-I response in BMDMs that was ablated in PARP14^−/−^ cells, demonstrating that PARP14 is required for the induction of IFN-I (10). To determine if PARP12 is also important for this IFN-I response we performed a similar experiment with PARP12^−/−^ BMDMs. Again, we observed an increase in IFN-*β* mRNA from N1347A as compared to WT virus infected PARP12^+/+^ BMDMs (Fig. 3). However, as opposed to results with PARP14^−/−^ BMDMs, we observed a similar increase of IFN-I mRNA during an N1347A infection in PARP12^−/−^ BMDMs, indicating that PARP12 is not required for IFN-I mRNA induction during a N1347A infection. We also looked at the mRNA levels of several key cytokines including IL-1*β*, IL-6, TNF-*α*, and CXCL-10 (Fig. 3). Again, there was no significant difference in mRNA levels between N1347A infected PARP12^+/+^ and PARP12^−/−^ BMDMs.

**Figure 3.**
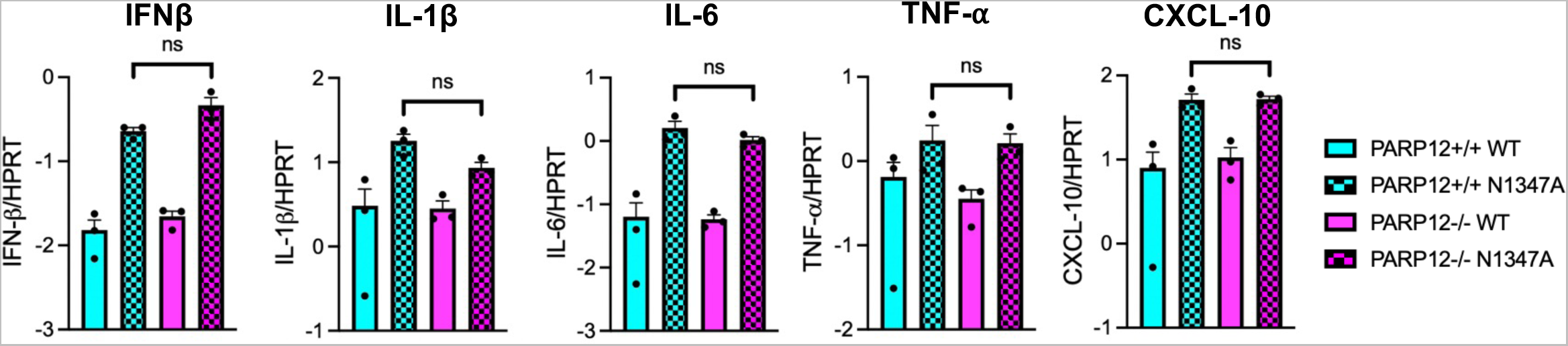
PARP12 does not contribute to enhance IFN and cytokine levels following BMDM infection with JHMV N1347A. PARP12+/+ and PARP12-/- BMDMs were infected with WT or N1347A JHM. Cells were collected in Trizol at 12 hpi and RNA levels were determined using RT-qPCR with specific primers for each gene of interest and normalized to HPRT. These data are from one experiment representative of two independent experiments (n=3).

### PARP12 deletion increases JHM N1347A replication following intracranial, but not intranasal infection

Despite multiple reports demonstrating PARP12 antiviral activity in cell culture, its ability to restrict virus replication *in vivo* has not been tested. Based on the increased replication seen in PARP12^−/−^ M0 macrophages, we hypothesized that JHMV N1347A replication in brains would be enhanced in PARP12^−/−^ mice. An intranasal infection with JHMV typically results in the infection of olfactory neurons and transneuronal spread via the olfactory bulb (OB) to primary, secondary, and tertiary connections of the OB. PARP12^+/+^ and PARP12^−/−^ mice were infected intranasally with 10^4^ PFU of JHMV WT or JHMV N1347A. Brains were then harvested at peak titer (5 dpi) (Fig. 4A). Similar to the BMDM titers and prior results (2), N1347A had reduced viral loads of approximately one-log in the PARP12^+/+^ mice compared to the WT virus, but surprisingly, there was no enhancement of N1347A replication in the PARP12^−/−^ mice (Fig. 4A). In addition, we performed immunohistochemistry staining of the forebrains of mice at 5 days-post-infection to assess virus spread (Fig. 4B, Fig. S4). PARP12^+/+^ and PARP12^−/−^ mice infected with the WT virus had extensive viral N-protein staining, in contrast, both PARP12^+/+^ and PARP12^−/−^ mice infected with N1347A had minimal N protein accumulation, indicating N1347A replicates poorly in the brain and that it is not restricted by PARP12 (Fig. 4B). We also looked at survival rates and weight loss following an intranasal infection. Both PARP12^+/+^ mice and PARP12^−/−^ mice succumbed to the WT virus at a similar rate with weight loss ranging from 10-15%. Following infection with N1347A, there was similar survival between the PARP12^+/+^ mice and the PARP12^−/−^ mice and little to no weight loss (Fig. 4C and D). We conclude that either a separate PARP or other innate immune factors act to suppress JHMV N1347A replication in the brains of PARP12^−/−^ mice following an intranasal infection.

**Figure 4:**
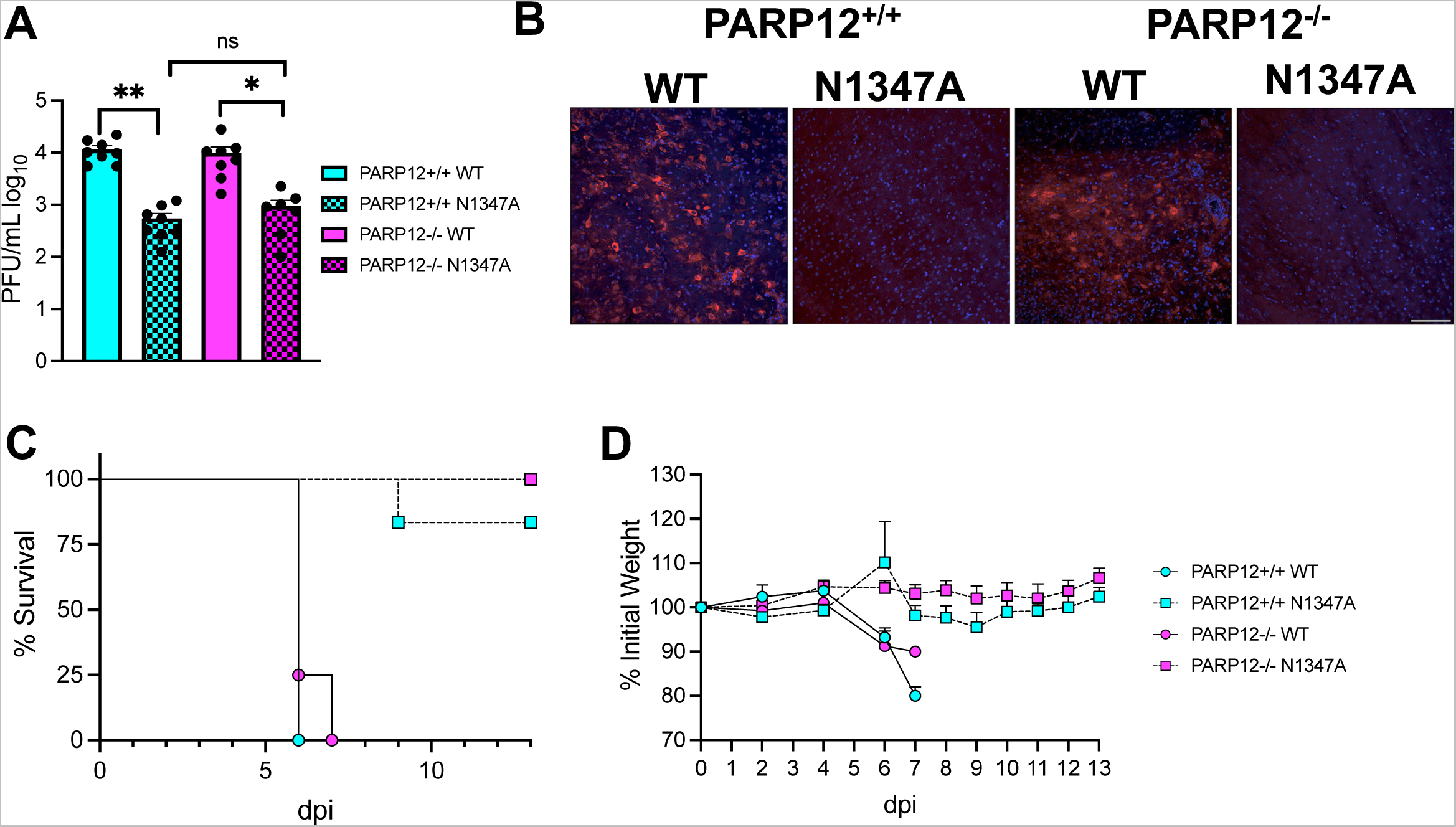
PARP12 is not required for the restriction of JHM virulence and replication following an IN infection. (A) PARP12^+/+^ mice and PARP12^−/−^ mice were infected with 10^4^ PFU of JHMV WT or N1347A virus via intranasal (IN) infection. Brains were harvested at 5 dpi and titers determined via plaque assay. The data in A is the combined results of 3 independent experiments (n=7-8 per group). (B) Infected brains were harvested at 5 dpi then forebrain sections were stained for MHV nucleocapsid (N) protein by IHC. n=2-4. (C-D) PARP12^+/+^ mice and PARP12^−/−^ mice were infected as described above. Survival and weight loss was monitored for 12 days (n=4-9 per group).

To test whether the route of infection into the brain would impact the ability of PARP12 to inhibit virus replication or reduce pathogenesis, we infected both PARP12^+/+^ mice and PARP12^−/−^ mice intracranially with 750 PFU of JHMV WT and JHMV N1347A and virus replication was measured at 4 dpi (peak titer). In the PARP12^+/+^ mice, there was a significant reduction in the replication of the JHM N1347A virus compared to the WT virus. However, in the PARP12^−/−^ mice there were several mice where N1347A replicated to the same level as WT virus, and there was no significant difference in the viral loads between WT and N1347A viruses, indicating that PARP12 at least partially restricted N1347A replication in the brains of mice (Fig. 5A). We then tested if there was an increase in disease in PARP12^−/−^ mice following an intracranial infection with N1347A. The WT virus caused 15-20% weight loss and 100% lethality in both PARP12^+/+^ mice and PARP12^−/−^ mice. However, following an intracranial N1347A infection, PARP12^+/+^ mice exhibited 10-15% weight loss and ~75% of infected mice survived. PARP12^−/−^ mice infected with N1347A also had 10-15% weight loss but only ~50% of infected mice survived, indicating a slight increase in lethality of PARP12^−/−^ mice following infection with N1347A, though this result was not statistically significant (Fig. 5B and C). These results demonstrate that the loss of PARP12 can in some cases impact the outcome of infection but continues to indicate that multiple PARPs or other innate immune factors contribute to reducing the instance of severe encephalitis following infection with N1347A.

**Figure 5.**
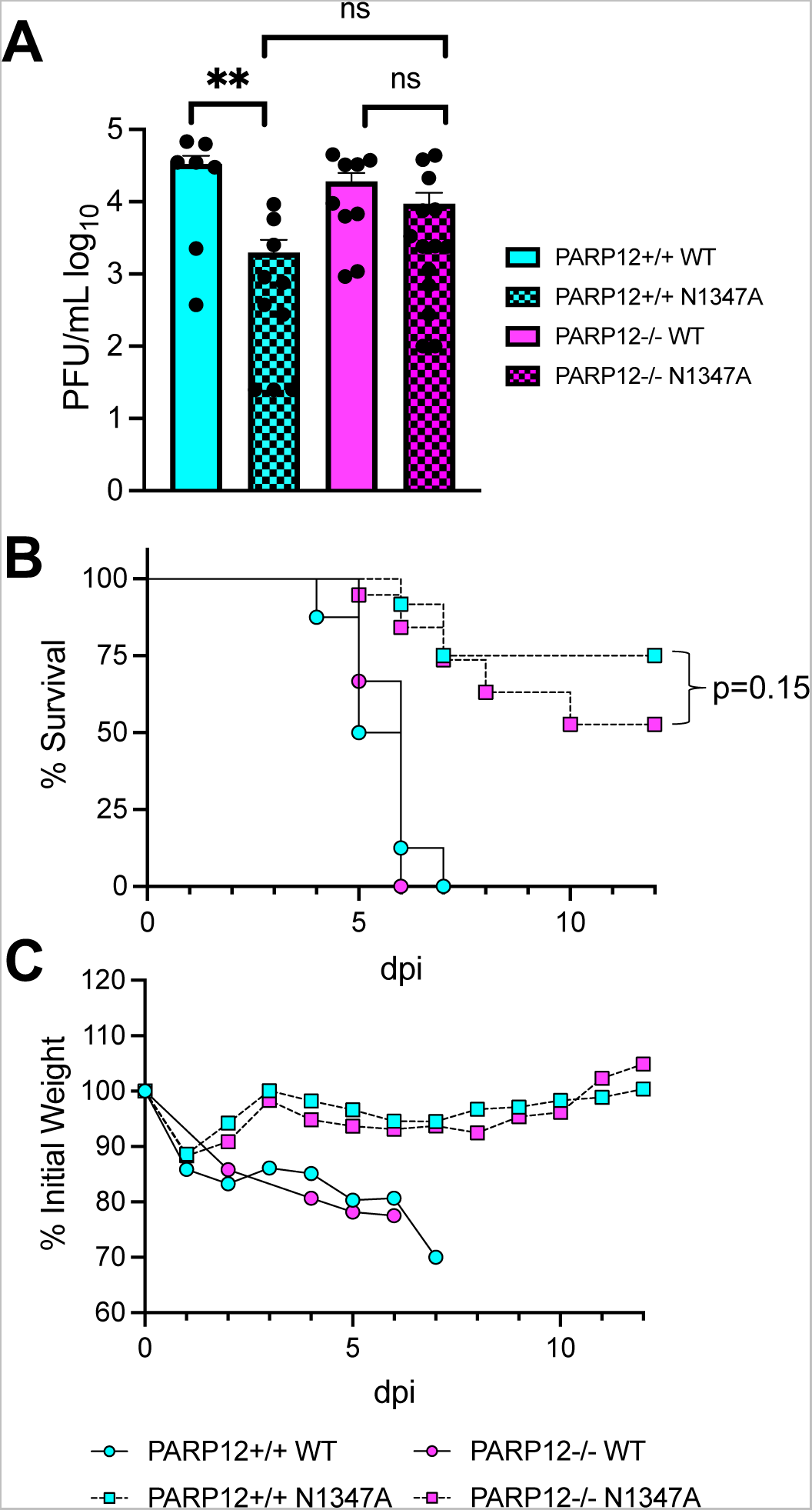
PARP12 KO mildly increases virus replication and lethality in the brain following an IC infection with N1347A. (A) PARP12^+/+^ mice and PARP12^−/−^ mice were infected IC with 750 PFU of MHV JHM WT or N1347A virus. (A) Brains were harvested at 4 dpi and titers were determined via plaque assay. Data in A is combined from >3 independent experiments (n=7-14 per group). (B and C) Mice were infected as described in A and survival and weight loss were monitored for 12 days (n=6-22 per group).

### PARP12 deletion enhances virus replication and pathology following infection of A59 N1348A in the liver

Given that MHV-A59 N1348A replication was rescued in PARP12^−/−^ BMDMs (Fig. 2A-B) and that PARP12 is well-expressed in the liver (Fig. 1D), we hypothesized that N1348A replication and pathogenesis would be enhanced in the livers of infected PARP12^−/−^ mice. To test this hypothesis, we infected 8-week-old mice with 500 PFU of A59 WT and N1348A viruses and measured virus replication at 3 dpi, the time of peak replication in the liver (34). While the viral loads of N1348A were significantly reduced compared to WT virus in PARP12^+/+^ mice, the viral loads of N1348A were rescued to WT levels in PARP12^−/−^ mice, indicating that PARP12 was indeed required to inhibit the replication of N1348A in livers (Fig. 6A). We next tested if there was increased disease in PARP12^−/−^ mice infected with N1348A compared to PARP12^+/+^ mice. Here we infected mice with 50,000 PFU of WT and N1348A to enable the development of clinical disease. In PARP12^+/+^ mice, WT virus caused 10-15% weight loss and ~50% of infected mice succumbed to the infection, whereas N1348A infected mice caused only mild weight loss and only one of the infected mice succumbed to infection. Even though viral loads of N1348A were rescued in PARP12^−/−^ mice, we were unable to detect any significant differences in the weight loss or survival of N1348A infected PARP12^−/−^ mice compared to the PARP12^+/+^ mice (Fig. 6B and C). To determine if PARP12 affects liver pathology in this animal model of MHV infection, we performed H&E staining on livers at the end point of infection. Livers were analyzed blindly and scored for inflammation, necrosis, and edema/fibrin depositions. Both PARP12^−/−^ and PARP12^+/+^ mice demonstrated substantial tissue damage following infection with WT virus, though about half of these mice still only scored a 1 or 0 in each category indicating a bimodal distribution (Fig. 7A-B). Following infection with N1348A, PARP12^+/+^ livers appeared largely normal, as only one of the PARP12^+/+^ mice infected with N1348A scored higher than a “1” in any category. In contrast, PARP12^−/−^ mice showed signs of tissue damage in their livers following N1348A infection, with several mice scoring a 2 or 3 in all 3 categories. These results indicate that PARP12 plays at least a part in preventing liver pathology following infection with a Mac1-mutant coronavirus.

**Figure 6.**
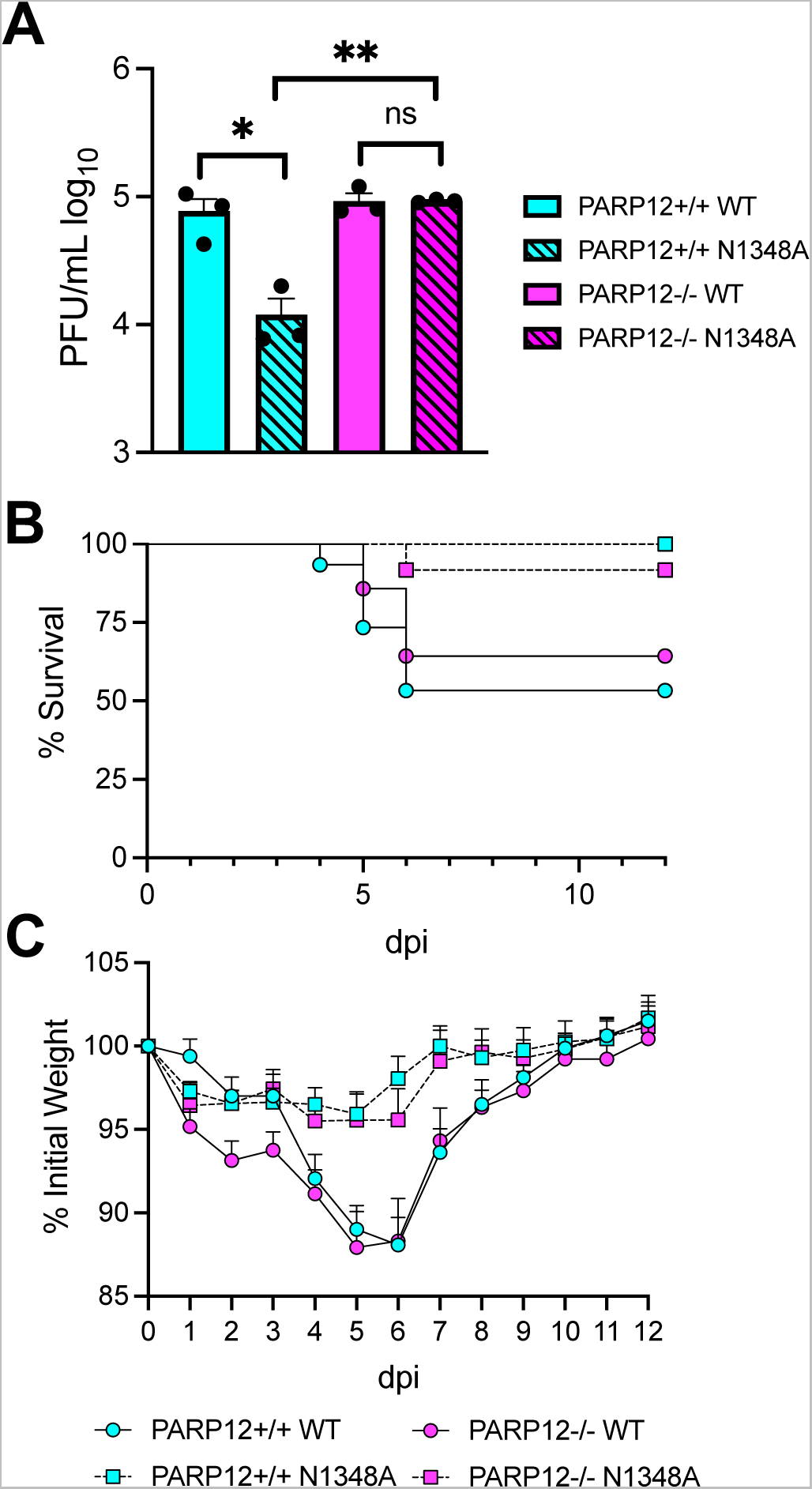
PARP12 KO increases N1348A replication in livers but does not impact survival or weight loss. (A) PARP12^+/+^ and PARP12^−/−^ mice were infected IP with 500 PFU of MHV-A59 WT or N1348A virus. Brains were harvested at 4 dpi and titers were determined via plaque assay. The results in A are from one experiment representative of 3 independent experiments (n=3 per group). (B-C) PARP12^+/+^ and PARP12^−/−^ mice were infected IP with 5×10^4^ PFU of MHV-A59 WT or N1348A virus and survival and weight loss was monitored for 12 days (n=12-16 per group).

**Fig. 7.**
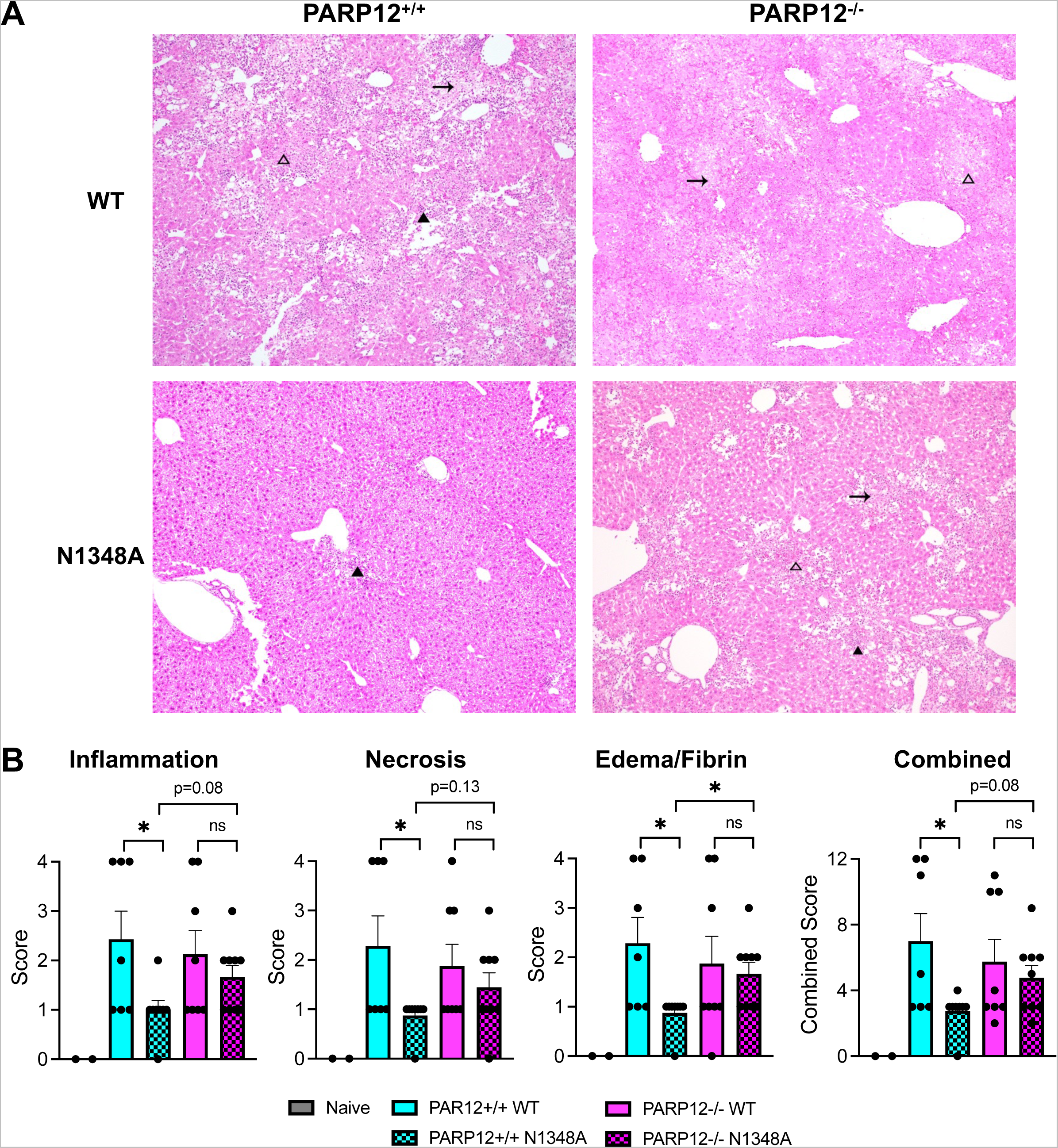
PARP12 is required to prevent severe liver pathology following A59 N1348A infection. (A) PARP12^+/+^ and PARP12^−/−^ mice were infected with 5×10^4^ PFU of MHV A59 WT or N1348A virus via IP injection and livers were harvested at 12 days post infection (dpi) and histological analysis of livers was performed (n=7-9 per group). Dark arrowheads represent inflammation, blank arrowheads represent necrosis, and arrows represent edema/fibrin. (B) Liver pathology was scored based on inflammation, necrosis, and edema/fibrin on a scale of 0-4 (see Methods). The combined score represents the combined scores for all 3 categories.

In total, we have found that PARP12 restricts the replication of a Mac1 mutant coronavirus in a cell culture and in mice in a cell type and tissue specific manner.

## DISCUSSION

ADP-ribosylation is an important yet under-recognized protein modification that plays numerous roles in cell biology. In recent years the importance of ADP-ribosylation in the context of virus infection has gotten increased attention as multiple studies across several positive-strand RNA virus families have indicated important roles for viral macrodomains in infection (35, 36). All CoVs encode for a macrodomain in nsp3 which can both bind to ADP-ribosylated proteins and reverse ADP-ribosylation via its enzyme activity (5–7). Work from our lab and others have demonstrated that this enzyme is critical for viral replication and pathogenesis and antagonizes IFN-I responses (2, 9, 10, 34). This includes SARS-CoV-2, as we recently found that Mac1-deleted SARS-CoV-2 is extremely attenuated in mice and induces a robust IFN response (37). However, many details of how ADP-ribosylation mediates these anti-viral effects remain unclear, including: *i)* what PARP(s) inhibit virus replication and pathogenesis and promote IFN-I responses, *ii)* how does ADP-ribosylation impact the virus lifecycle, and *iii)* how do these PARPs mechanistically inhibit virus replication.

Several PARPs are induced by interferon, including PARP7, PARP9, PARP10, PARP11, PARP12, PARP13, and PARP14. The first indication that the interferon-induced ARTDs may restrict N1347A replication was the finding that N1347A replicates to near WT virus levels in IFNAR^−/−^ cells. A subsequent siRNA screen of IFN-stimulated PARPs found that the knockdown of PARP12 and, to a lesser extent, PARP14 could enhance N1347A replication (10). To expand upon our siRNA results, we developed PARP12 knockout mice to confirm our siRNA results and define the role of PARP12 during an *in vivo* virus infection. While PARP12 can inhibit the replication of several families of viruses, including alphaviruses, flaviviruses, and rhabdoviruses (18, 23, 24), no one has demonstrated a role for this host factor *in vivo*.

One of the first notable findings here was that, despite being an ISG, PARP12 mRNA is relatively well expressed in several tissues of WT mice, especially the heart, lung, liver, and even the testes of male mice. However, the lack of a reliable antibody has prevented us from examining PARP12 expression at the protein level in tissue and even in cell culture. PARPs exist at low levels in cells which are difficult to detect even by mass spec (38, 39). Thus, novel detection methods may be required to identify PARP12 protein expression in cells and tissue.

Despite difficulties in detecting PARP12 protein, we demonstrated that the deletion of the PARP12 gene fully or partially rescued the replication of a Mac1 mutant MHV, but not WT MHV, in both in cell culture and in mice. These results demonstrate that PARP12 can function to inhibit CoV replication, but also that its function is effectively thwarted by Mac1. The mechanism by which PARP12 represses coronavirus replication remains unknown. PARP12 has been shown to relocate to stress granules from the Golgi following induction of cell stress, where it could potentially function to inhibit viral protein translation (18, 26). However, the very low abundance of PARP12 and lack of effective antibodies for PARP12 have limited our ability to detect its localization during infection. When expressed from a VEEV vector, PARP12 repressed the VEEV replication and expression of nsp2, a marker for general protein expression. When further investigated, PARP12 expression led to decreased cellular translation through interactions with ribosomes, specifically polysomes (18). PARP12 catalytic activity was largely required for its ability to inhibit translational inhibition, but only played a small role in its antiviral activity, making it unclear exactly how PARP12 represses VEEV replication (40). PARP12 contains several Zn-Finger and WWE domains like PARP7 and PARP13 (Zinc-antiviral protein or ZAP). ZAP has several known antiviral activities, including blocking protein translation, degrading viral RNAs, and others, though it lacks ADP-ribosyltransferase activity due to the mutation of several key enzymatic residues (17). One hypothesis is that PARP12 positively regulates itself by auto-MARylation which activates its additional domains to restrict CoV replication. Alternatively, it could ADP-ribosylate other host or viral proteins which could activate their functions to repress virus replication. We previously determined that the CoV N protein is ADP-ribosylated; however, the level of ADP-ribosylation was unchanged in the N1347A mutant virus infection, indicating that the ADP-ribosylation of N protein does not likely impact the phenotypes described in this or previous reports (41). Further investigation into the mechanisms of PARP12’s antiviral activity and its impact on the viral lifecycle are needed to fully uncover the basis for its antiviral activity.

PARP12 is likely not the only PARP that inhibits Mac1 mutant MHV. Our results revealed a substantial level of both cell-type and tissue specific activity for PARP12. Our observation that PARP12 knockout could enhance both JHMV-N1347A and A59-N1348A viruses to WT levels in BMDMs, but has no impact on their replication in BMDCs, demonstrates that other PARPs must have redundant functions. The idea that PARPs may be redundant is not without precedent. For instance, PARP1 or PARP2 KO mice are developmentally normal, but a double knockout is embryonic lethal, and PARP1 shares many similar functions with other nuclear PARPs (29, 42, 43). Furthermore, PARP5a/5b share multiple functions and recently were found to target MAVS for PARylation and subsequent proteasome degradation (44, 45). In addition to redundancy, our results indicate that multiple PARPs likely work together to fully attenuate Mac1 mutant virus. For instance, in the livers, the loss of PARP12 enhanced virus replication and increased virally induced pathology, but the N1348A virus still did not cause significant weight loss or lethality in infected mice, indicating a role for additional PARPs in driving disease phenotypes.

These results drive the question, what other PARP may have redundant function or cooperate with PARP12 to fully attenuate a Mac1 mutant virus? Expression of PARP7 and PARP10 blocked cellular translation and VEEV replication to nearly identical levels as PARP12, indicating similar function (40). PARP12 was shown to interact with PARP14 (46), and our previous study found that knockdown of PARP14 also slightly enhanced N1347A virus replication in BMDMs. Finally, PARP11 was shown to function along with PARP12 to restrict Zika virus infection by targeting NS1 and NS3 for degradation (25). PARP10, PARP11, and PARP14 do not share many of the same domains as PARP12, thus, it is not clear which PARP may be functionally redundant with PARP12 or provide additional functions to prevent viral pathogenesis. We are actively pursuing which additional PARPs might contribute to the attenuation of Mac1 mutant coronaviruses.

In total, these results have revealed extensive new insight into the role of the PARP12 protein in the antiviral response to coronavirus infection both in cell culture and in mice. Understanding the interactions between Mac1 and ARTs could have important implications in coronavirus evolution and antiviral drug and vaccine development.

## METHODS

### Cell culture

HeLa cells expressing the MHV receptor carcinoembryonic antigen-related cell adhesion molecule 1 (CEACAM1) (HeLa-MHVR) were grown in Dulbecco’s Modified Eagle Medium (DMEM) supplemented with 10% fetal bovine serum (FBS), 100 U/ml penicillin and 100 mg/ml streptomycin, HEPES, sodium pyruvate, non-essential amino acids, and L-glutamine. Bone marrow-derived macrophages (BMDMs) sourced from PARP12^+/+^ and PARP12^−/−^ mice were differentiated by incubating cells in Roswell Park Memorial Institute (RPMI) media supplemented with 10% L929 cell supernatants (unless otherwise stated), 10% FBS, sodium pyruvate, 100 U/ml penicillin and 100 mg/ml streptomycin, and L-glutamine for seven days. Bone marrow derived dendritic cells (BMDCs) were differentiated by incubating cells with Roswell Park Memorial Institute (RPMI) media supplemented with 10% FBS, sodium pyruvate, 100 U/ml penicillin and 100 mg/ml streptomycin, L-glutamine, and 20 ng/ml GM-CSF for seven days. All cells were washed and replaced with fresh media every day after the 4^th^ day.

### Mice

C57BL/6N-*Parp12^tm1.1(KOMP)Vlcg^*/MbpMmucd (PARP12^−/−^) was produced by the Mouse Biology Program using ES cell clone 15401A-D1, which was provided to KOMP by Velocigene-Regeneron. After microinjection and germline transmission, mice that contained the reporter-tagged null allele (tm1) were bred to Cre-expressing mice. This resulted in removal of the *β*-actin promoter and the Neomycin gene it activated (tm1.1). The tm1.1 allele remains a lacZ reporter and is a non-conditional knock-out of the gene (Fig. 1A). Please see the following link for targeting strategy information and images: https://www.mousephenotype.org/understand/the-data/allele-design/. All animal procedures were conducted according to the Transgenic and Gene-Targeting Facility’s and the Fehr lab Animal Care and Use Protocol approved by the KUMC and KU Institutional Animal Care and Use Committees (IACUC), respectively, following guidelines set forth in the Guide for the Care and Use of Laboratory Animals. The KUMC transgenic mouse facility performed *in vitro* fertilization of the sperm from PARP12^+/−^ mice with pathogen-free C57BL/6NJ (B6) mice to reestablish the mouse line. Pathogen-free C57BL/6NJ (B6) mice were originally purchased from Jackson Laboratories. prepubertal female mice were superovulated by i.p. administration of 5 IU P.G. 600 (PMSG)(Intervet Inc.), followed 48 hours later by i.p. administration of 5 IU human chorionic gonadotropin (hCG)(Sigma, #C1063). The next morning, females were euthanized 15 hours after administration of hCG for the collection of oviducts. Cumulus-oocyte complexes were released from the oviducts under oil and dragged into a 90ml drop of CARD medium (COSMO BIO USA, kit KYD-005-EX) and incubated at 37°C, 6% CO_2_, 5% O_2_ for 30 minutes to one hour. A straw of frozen PARP12+/- sperm was removed from liquid nitrogen, held in air for five seconds, and submerged in a 37°C water bath for 10 minutes. The sperm sample was expelled into a 90ml drop of CARD Preincubation Medium and incubated. After 30 minutes, a 10ml aliquot of the sperm suspension was withdrawn from the preincubation drop and released into the CARD drop containing the oocytes. Gametes were co-incubated for four hours, at which time the oocytes were washed free of the sperm and moved to a drop of KSOM culture medium (Millipore Sigma, #MR-101-D) for overnight culture. Fertilized oocytes were scored and separated the next morning at the two-cell stage for surgical transfer to pseudopregnant CD-1 recipient females (Charles River, #022). Heterozygote mice were transferred to the University of Kansas Animal Care Unit and heterozygote pairs were bred to create PARP12^+/+^, PARP12^+/−^, and PARP12^−/−^ mice. Mice were genotyped using primers F 5’-TGTGGGTGTATTTTCACACAAGC-3’ and R 5’-TGTACCACTGGAGAAGGATGAAGCC-3’ to detect the PARP12 WT allele (224 bp) and primers F 5’-AAAAGCAAACTGGACCACAAGACCC-3’ and R 5’-ACTTGCTTTAAAAAACCTCCCACA-3’ to detect the PARP12 KO allele (950 bp).

### Virus infection

Recombinant MHV-JHMV was previously described (2) and recombinant MHV-A59 was kindly provided by Dr. Susan Weiss. Cells were infected with recombinant MHV at a multiplicity of infection (MOI) of 0.05-0.1 PFU/cell with a 60 min adsorption phase. For MHV-A59 *in vivo* infections, 8-12 week-old male and female mice were inoculated via an intraperitoneal injection with either 500 or 5×10^4^ PFU of recombinant A59 in a total volume of 200µl PBS. For JHMV *in vivo* infections, 5-8 week-old male and female mice were anesthetized with ketamine/xylazine and inoculated intranasally with either 1×10^4^ PFU recombinant JHMV in a total volume of 12 μl DMEM, or 5-6 week old male and female mice were anesthetized with ketamine/xylazine and inoculated intracranially with 750 PFU of recombinant JHMV in a total volume of 30 µl DMEM. To obtain viral titers from infected animals, mice were sacrificed, and brain tissue was collected and homogenized in DMEM. Viral titers were determined by plaque assay using HeLa-MHVR cells.

### Real-time qPCR analysis

RNA was isolated from BMDMs using TRIzol (Invitrogen) and cDNA was prepared using MMLV-reverse transcriptase as per manufacturer’s instructions (Thermo Fisher Scientific). Quantitative real-time PCR (qRT-PCR) was performed on a QuantStudio3 real-time PCR system using PowerUp SYBR Green Master Mix (Thermo Fisher Scientific). Primers used for qPCR are listed in Table S2. Cycle threshold (C_T_) values were normalized to the housekeeping gene hypoxanthine phosphoribosyltransferase (HPRT) by the following equation: C_T_ = C_T(gene of interest)_ - C_T(HPRT)_. Results are shown as a ratio to HPRT calculated as 2^−ΔCT^.

### Immunohistochemistry Staining

Mice were perfused intracardially with 4% formaldehyde (FA) diluted in 1X HBSS. After perfusion, each mouse brain was dissected and immersed in fresh 4% FA in individual tubes for post-fixation. The forebrain region was cut, placed on a freezing stage at −20°C and sectioned rostral-to-caudal at 30 µm in intervals (skip 60 µm between sections) using a sliding block microtome (American Optical Spencer 860 with Cryo-Histomat MK-2 controller). Four to six forebrain sections per group were moved individually on a 24-well plate using a camel hairbrush < 1.59 mm (Electron Microscopy Sciences, Cat. No. 65575-02), free-floating on HBSS/0.1% sucrose (HBSS/Su) at room temperature (RT) for rinsing. To perform single immunohistochemistry, forebrain sections were permeabilized with HBSS/Su + 0.1% saponin (HBSS/Su/Sap; 2x, 5 min each), blocked with HBSS/Su/Sap + 0.1% Triton X-100 + 3% rabbit serum (1x, 1hr at RT), rinsed and incubated with the primary antibody mouse anti-N (1:5000) diluted in HBSS/Su/Sap + 3% rabbit serum O/N at 4 °C. The next day, forebrain sections were rinsed with HBSS/Su/Sap (3x, 5 min each) and incubated with the secondary antibody Alexa Fluor 594 rabbit anti-mouse (1:200) diluted in HBSS/Su/Sap (3 hrs at RT), rinsed with HBSS/Su/Sap (2x, 5 min each), HBSS/Su (1x, 5 min) and HBSS (1x, 5 min). DAPI (10 µM) diluted in HBSS was added for nuclear counterstain (1hr at RT) and then rinsed with HBSS (2x, 5 min each). Forebrain sections were carefully moved on a microscope slide using a camel hairbrush < 1.59 mm, mounted with Vectashield® Antifade mounting medium, and cover-slipped (22 x 50 cover glass; No. 1.5 thickness) for imaging.

### Image acquisition

Fluorescent images were acquired using a TCS SPE Laser Scanning Confocal Upright Microscope (Leica Microsystems, DM6-Q model), with the 405 nm and 561 nm laser lines, an Olympus 20X/0.75NA UPlanSApo infinity corrected, 8-bit spectral PMT detector and a Leica LAS X Imaging software (version 3.5.7.23225). Two to four images were taken per section. Anti-N + Alexa Fluor 594 signal was detected using 561 nm excitation (35% laser intensity), 600-620 nm emission range, 700 V PMT gain, and 0% offset, while DAPI signal was detected using 405 nm excitation (6% laser intensity), 430-480 nm emission range, 700 V PMT gain, and 0% offset. Images were captured at 1024 x 1024-pixel resolution with a scan speed at 400, no bidirectional scanning, a zoom factor at 1.0, Pinhole 1.0 AU = 75.54 µm (550 µm x 550 µm image size; 537.63 nm x 537.63 nm pixel size; 2.057 µm optical section and 0.69 µm step size). Leica LAS X software_3D Viewer was used for post-processing to create figure plates, while raw data was exported as .tiff for relative fluorescent data analysis. All the workflow design, sample preparation, processing and imaging was performed in the Microscopy and Analytical Imaging Resource Core Laboratory (RRID:SCR_021801) at The University of Kansas.

### H&E Staining

The livers were perfused and placed in 10% of formalin. The representative liver sections were then processed for hematoxylin and eosin (H&E) staining. The liver lesions were blindly scored by an American College of Veterinary Pathology Board-certified pathologist. The lesions were scored on a scale of 0-10% (score 1), 10-40% (score 2), 40-70% (score 3) and >70% (score 4) and cumulative scores were obtained for each mouse. The lesions scored were inflammation, necrosis, and edema/fibrin.

### Flow Cytometry

Mouse spleens were excised and placed in PBS. Samples were smashed into single cell suspension and filtered through a 40uM filter to create a single cell suspension. Single cell suspension was counted and resuspended to desired concentration (dependent on experiment) in PBS. Single cell suspensions were used for staining and flow cytometric analysis. Cell were stained in serum free PBS. All flow cytometry was completed on a spectral cytometer the Cytek Aurora with a 5 laser system (355nm, 405nm, 488nm, 561nm, 640nm). Single color stain OneComp eBeads (Thermo Fisher) were used for unmixing. Unmixed files were analyzed using FlowJo Software (BD Biosciences, San Diego, California). Antibodies used in various combinations (depending on experiment) are as follows: Ghost Viability Dye (v510, Tonbo Biosciences, 1:1000 dilution), CD45 (BUV395, BD Biosciences, 1:500 dilution, clone 30-F11), CD3 (PE-Cy5, Tonbo,1:200, clone 145-2C11), CD4 (BV605, Biolegend, 1:200, clone GK1.5),CD8a (APC-Cy7, Tonbo, 1:200, clone 53-6.7), CD11c (PE-Cy5.5, Tonbo, 1:100, clone N418), CD11b (PerCP-Cy5.5, Tonbo, 1:100, clone M1/70), CD19 (BV711, Biolegend, 1:400, clone 6D5), CD69 (PE, Biolegend, 1:200, clone H1-2F3), CD103 (PerCP-ef710, Thermo Fisher, 1:200, clone 2E7), CD44 (AlexaFluor700, Tonbo, 1:200, clone IM7), CD62L (PE-Cy7, Tonbo, 1:200, clone MEL-14), MHC II (I-A/I-E) (SuperBright645, Thermo Fisher, 1:200,M5/114.15.2), MHC I (H-2Kb/Db) (FITC, Biolegend, 1:200, clone 28.8-6), Ly6C (BV785, Biolegend, 1:300, clone HK1.4), Ly6G (PE-efluor610, company, 1:300, clone IA8), B220 (APC-Cy5.5, Thermo Fisher, 1:200, clone RA3-6B2)PDCA-1/CD317 (Pacific Blue, Biolegend, 1:200, clone 129C1), F4/80 (Pacific Orange, Thermo Fisher, 1:100, clone BM8). All surface markers were stained in PBS at 4°C in the dark. Samples were fixed in 1% PFA.

### Statistics

A Student’s *t* test was used to analyze differences in mean values between groups. All results are expressed as means ± standard errors of the means (SEM). Differences in survival were calculated using a Kaplan-Meier log-rank test. P values of ≤0.05 were considered statistically significant (*, P≤0.05; **, P≤0.01; ***, P≤0.001; ****, P ≤0.0001; n.s., not significant).

## ACKNOWLEDGEMENTS

We thank members of the Fehr and Davido laboratory at KU for valuable discussion. We thank Dr. Stanley Perlman (University of Iowa) for reagents and critical reading of the manuscript and Dr. Susan Weiss (University of Pennsylvania) for reagents. These studies were supported by the KU microscopy and analytical imaging (MAI) facility, and the Transgenic and Gene Targeting Shared Resource (TGTSR) (NIH Funding P30CA168524) of the University of Kansas Cancer Center and University of Kansas Medical Center. The mouse strain used for this research project, C57BL/6N-*Parp12^tm1.1(KOMP)Vlcg^*/MbpMmucd, RRID:MMRRC_048982-UCD, was obtained from the Mutant Mouse Resource and Research Center (MMRRC) at University of California at Davis, an NIH-funded strain repository, and was donated to the MMRRC by The KOMP Repository, University of California, Davis; Originating from Kent Lloyd, UC Davis Mouse Biology Program.

## REFERENCES

1. Perlman S. 2020. Another Decade, Another Coronavirus. N Engl J Med 382:760–762.

2. Fehr AR, Athmer J, Channappanavar R, Phillips JM, Meyerholz DK, Perlman S. 2015. The nsp3 macrodomain promotes virulence in mice with coronavirus-induced encephalitis. J Virol 89:1523–36.

3. Fehr AR, Perlman S. 2015. Coronaviruses: an overview of their replication and pathogenesis. Methods Mol Biol 1282:1–23.

4. Comar CE, Otter CJ, Pfannenstiel J, Doerger E, Renner DM, Tan LH, Perlman S, Cohen NA, Fehr AR, Weiss SR. 2022. MERS-CoV endoribonuclease and accessory proteins jointly evade host innate immunity during infection of lung and nasal epithelial cells. Proc Natl Acad Sci U S A 119:e2123208119.

5. Li C, Debing Y, Jankevicius G, Neyts J, Ahel I, Coutard B, Canard B. 2016. Viral Macro Domains Reverse Protein ADP-Ribosylation. J Virol 90:8478–86.

6. Egloff MP, Malet H, Putics A, Heinonen M, Dutartre H, Frangeul A, Gruez A, Campanacci V, Cambillau C, Ziebuhr J, Ahola T, Canard B. 2006. Structural and functional basis for ADP-ribose and poly(ADP-ribose) binding by viral macro domains. J Virol 80:8493–502.

7. Saikatendu KS, Joseph JS, Subramanian V, Clayton T, Griffith M, Moy K, Velasquez J, Neuman BW, Buchmeier MJ, Stevens RC, Kuhn P. 2005. Structural basis of severe acute respiratory syndrome coronavirus ADP-ribose-1’’-phosphate dephosphorylation by a conserved domain of nsP3. Structure 13:1665–75.

8. Fehr AR, Jankevicius G, Ahel I, Perlman S. 2018. Viral Macrodomains: Unique Mediators of Viral Replication and Pathogenesis. Trends Microbiol 26:598–610.

9. Fehr AR, Channappanavar R, Jankevicius G, Fett C, Zhao J, Athmer J, Meyerholz DK, Ahel I, Perlman S. 2016. The Conserved Coronavirus Macrodomain Promotes Virulence and Suppresses the Innate Immune Response during Severe Acute Respiratory Syndrome Coronavirus Infection. mBio 7.

10. Grunewald ME, Chen Y, Kuny C, Maejima T, Lease R, Ferraris D, Aikawa M, Sullivan CS, Perlman S, Fehr AR. 2019. The coronavirus macrodomain is required to prevent PARP-mediated inhibition of virus replication and enhancement of IFN expression. PLoS Pathog 15:e1007756.

11. Alhammad YMO, Fehr AR. 2020. The Viral Macrodomain Counters Host Antiviral ADP-Ribosylation. Viruses 12.

12. Gupte R, Liu Z, Kraus WL. 2017. PARPs and ADP-ribosylation: recent advances linking molecular functions to biological outcomes. Genes Dev 31:101–126.

13. Palazzo L, Mikolcevic P, Mikoc A, Ahel I. 2019. ADP-ribosylation signalling and human disease. Open Biol 9:190041.

14. Holbourn KP, Shone CC, Acharya KR. 2006. A family of killer toxins. Exploring the mechanism of ADP-ribosylating toxins. FEBS J 273:4579–93.

15. Hottiger MO, Hassa PO, Luscher B, Schuler H, Koch-Nolte F. 2010. Toward a unified nomenclature for mammalian ADP-ribosyltransferases. Trends Biochem Sci 35:208–19.

16. Luscher B, Ahel I, Altmeyer M, Ashworth A, Bai P, Chang P, Cohen M, Corda D, Dantzer F, Daugherty MD, Dawson TM, Dawson VL, Deindl S, Fehr AR, Feijs KLH, Filippov DV, Gagne JP, Grimaldi G, Guettler S, Hoch NC, Hottiger MO, Korn P, Kraus WL, Ladurner A, Lehtio L, Leung AKL, Lord CJ, Mangerich A, Matic I, Matthews J, Moldovan GL, Moss J, Natoli G, Nielsen ML, Niepel M, Nolte F, Pascal J, Paschal BM, Pawlowski K, Poirier GG, Smith S, Timinszky G, Wang ZQ, Yelamos J, Yu X, Zaja R, Ziegler M. 2021. ADP-ribosyltransferases, an update on function and nomenclature. FEBS J doi:10.1111/febs.16142.

17. Fehr AR, Singh SA, Kerr CM, Mukai S, Higashi H, Aikawa M. 2020. The impact of PARPs and ADP-ribosylation on inflammation and host-pathogen interactions. Genes Dev 34:341–359.

18. Atasheva S, Akhrymuk M, Frolova EI, Frolov I. 2012. New PARP gene with an anti-alphavirus function. J Virol 86:8147–60.

19. Yamada T, Horimoto H, Kameyama T, Hayakawa S, Yamato H, Dazai M, Takada A, Kida H, Bott D, Zhou AC, Hutin D, Watts TH, Asaka M, Matthews J, Takaoka A. 2016. Constitutive aryl hydrocarbon receptor signaling constrains type I interferon-mediated antiviral innate defense. Nat Immunol 17:687–94.

20. Wang F, Zhao M, Chang B, Zhou Y, Wu X, Ma M, Liu S, Cao Y, Zheng M, Dang Y, Xu J, Chen L, Liu T, Tang F, Ren Y, Xu Z, Mao Z, Huang K, Luo M, Li J, Liu H, Ge B. 2022. Cytoplasmic PARP1 links the genome instability to the inhibition of antiviral immunity through PARylating cGAS. Mol Cell 82:2032–2049 e7.

21. Zhang W, Guo J, Chen Q. 2022. Role of PARP-1 in Human Cytomegalovirus Infection and Functional Partners Encoded by This Virus. Viruses 14.

22. Guo T, Zuo Y, Qian L, Liu J, Yuan Y, Xu K, Miao Y, Feng Q, Chen X, Jin L, Zhang L, Dong C, Xiong S, Zheng H. 2019. ADP-ribosyltransferase PARP11 modulates the interferon antiviral response by mono-ADP-ribosylating the ubiquitin E3 ligase beta-TrCP. Nat Microbiol 4:1872–1884.

23. Liu SY, Sanchez DJ, Aliyari R, Lu S, Cheng G. 2012. Systematic identification of type I and type II interferon-induced antiviral factors. Proc Natl Acad Sci U S A 109:4239–44.

24. Li L, Zhao H, Liu P, Li C, Quanquin N, Ji X, Sun N, Du P, Qin CF, Lu N, Cheng G. 2018. PARP12 suppresses Zika virus infection through PARP-dependent degradation of NS1 and NS3 viral proteins. Sci Signal 11.

25. Li L, Shi Y, Li S, Liu J, Zu S, Xu X, Gao M, Sun N, Pan C, Peng L, Yang H, Cheng G. 2021. ADP-ribosyltransferase PARP11 suppresses Zika virus in synergy with PARP12. Cell Biosci 11:116.

26. Welsby I, Hutin D, Gueydan C, Kruys V, Rongvaux A, Leo O. 2014. PARP12, an interferon-stimulated gene involved in the control of protein translation and inflammation. J Biol Chem 289:26642–26657.

27. Heer CD, Sanderson DJ, Voth LS, Alhammad YMO, Schmidt MS, Trammell SAJ, Perlman S, Cohen MS, Fehr AR, Brenner C. 2020. Coronavirus infection and PARP expression dysregulate the NAD metabolome: An actionable component of innate immunity. J Biol Chem 295:17986–17996.

28. Lee S, Lee YS, Choi Y, Son A, Park Y, Lee KM, Kim J, Kim JS, Kim VN. 2021. The SARS-CoV-2 RNA interactome. Mol Cell 81:2838–2850 e6.

29. Kelleher AM, Setlem R, Dantzer F, DeMayo FJ, Lydon JP, Kraus WL. 2021. Deficiency of PARP-1 and PARP-2 in the mouse uterus results in decidualization failure and pregnancy loss. Proc Natl Acad Sci U S A 118.

30. Celik-Ozenci C, Tasatargil A. 2013. Role of poly(ADP-ribose) polymerases in male reproduction. Spermatogenesis 3:e24194.

31. Meyer-Ficca ML, Ihara M, Bader JJ, Leu NA, Beneke S, Meyer RG. 2015. Spermatid head elongation with normal nuclear shaping requires ADP-ribosyltransferase PARP11 (ARTD11) in mice. Biol Reprod 92:80.

32. Osada T, Ogino H, Hino T, Ichinose S, Nakamura K, Omori A, Noce T, Masutani M. 2010. PolyADP-ribosylation is required for pronuclear fusion during postfertilization in mice. PLoS One 5.

33. Voth LS, O’Connor JJ, Kerr CM, Doerger E, Schwarting N, Sperstad P, Johnson DK, Fehr AR. 2021. Unique Mutations in the Murine Hepatitis Virus Macrodomain Differentially Attenuate Virus Replication, Indicating Multiple Roles for the Macrodomain in Coronavirus Replication. J Virol 95:e0076621.

34. Eriksson KK, Cervantes-Barragan L, Ludewig B, Thiel V. 2008. Mouse hepatitis virus liver pathology is dependent on ADP-ribose-1’’-phosphatase, a viral function conserved in the alpha-like supergroup. J Virol 82:12325–34.

35. McPherson RL, Abraham R, Sreekumar E, Ong SE, Cheng SJ, Baxter VK, Kistemaker HA, Filippov DV, Griffin DE, Leung AK. 2017. ADP-ribosylhydrolase activity of Chikungunya virus macrodomain is critical for virus replication and virulence. Proc Natl Acad Sci U S A 114:1666–1671.

36. Parvez MK. 2015. The hepatitis E virus ORF1 ‘X-domain’ residues form a putative macrodomain protein/Appr-1’’-pase catalytic-site, critical for viral RNA replication. Gene 566:47–53.

37. Alhammad YM, Parthasarathy S, Ghimire R, O’Connor JJ, Kerr CM, Pfannenstiel JJ, Chanda D, Miller CA, Unckless RL, Zuniga S, Enjuanes L, More S, Channappanavar R, Fehr AR. 2023. SARS-CoV-2 Mac1 is required for IFN antagonism and efficient virus replication in mice. bioRxiv doi:10.1101/2023.04.06.535927:2023.04.06.535927.

38. Bouhaddou M, Memon D, Meyer B, White KM, Rezelj VV, Correa Marrero M, Polacco BJ, Melnyk JE, Ulferts S, Kaake RM, Batra J, Richards AL, Stevenson E, Gordon DE, Rojc A, Obernier K, Fabius JM, Soucheray M, Miorin L, Moreno E, Koh C, Tran QD, Hardy A, Robinot R, Vallet T, Nilsson-Payant BE, Hernandez-Armenta C, Dunham A, Weigang S, Knerr J, Modak M, Quintero D, Zhou Y, Dugourd A, Valdeolivas A, Patil T, Li Q, Huttenhain R, Cakir M, Muralidharan M, Kim M, Jang G, Tutuncuoglu B, Hiatt J, Guo JZ, Xu J, Bouhaddou S, Mathy CJP, Gaulton A, Manners EJ, et al. 2020. The Global Phosphorylation Landscape of SARS-CoV-2 Infection. Cell 182:685–712 e19.

39. Stukalov A, Girault V, Grass V, Karayel O, Bergant V, Urban C, Haas DA, Huang Y, Oubraham L, Wang A, Hamad MS, Piras A, Hansen FM, Tanzer MC, Paron I, Zinzula L, Engleitner T, Reinecke M, Lavacca TM, Ehmann R, Wolfel R, Jores J, Kuster B, Protzer U, Rad R, Ziebuhr J, Thiel V, Scaturro P, Mann M, Pichlmair A. 2021. Multilevel proteomics reveals host perturbations by SARS-CoV-2 and SARS-CoV. Nature 594:246–252.

40. Atasheva S, Frolova EI, Frolov I. 2014. Interferon-stimulated poly(ADP-Ribose) polymerases are potent inhibitors of cellular translation and virus replication. J Virol 88:2116–30.

41. Grunewald ME, Fehr AR, Athmer J, Perlman S. 2018. The coronavirus nucleocapsid protein is ADP-ribosylated. Virology 517:62–68.

42. Menissier de Murcia J, Ricoul M, Tartier L, Niedergang C, Huber A, Dantzer F, Schreiber V, Ame JC, Dierich A, LeMeur M, Sabatier L, Chambon P, de Murcia G. 2003. Functional interaction between PARP-1 and PARP-2 in chromosome stability and embryonic development in mouse. EMBO J 22:2255–63.

43. Wang ZQ, Auer B, Stingl L, Berghammer H, Haidacher D, Schweiger M, Wagner EF. 1995. Mice lacking ADPRT and poly(ADP-ribosyl)ation develop normally but are susceptible to skin disease. Genes Dev 9:509–20.

44. Damale MG, Pathan SK, Shinde DB, Patil RH, Arote RB, Sangshetti JN. 2020. Insights of tankyrases: A novel target for drug discovery. Eur J Med Chem 207:112712.

45. Xu YR, Shi ML, Zhang Y, Kong N, Wang C, Xiao YF, Du SS, Zhu QY, Lei CQ. 2022. Tankyrases inhibit innate antiviral response by PARylating VISA/MAVS and priming it for RNF146-mediated ubiquitination and degradation. Proc Natl Acad Sci U S A 119:e2122805119.

46. Caprara G, Prosperini E, Piccolo V, Sigismondo G, Melacarne A, Cuomo A, Boothby M, Rescigno M, Bonaldi T, Natoli G. 2018. PARP14 Controls the Nuclear Accumulation of a Subset of Type I IFN-Inducible Proteins. J Immunol 200:2439–2454.

